# Fiberoptic Hemodynamic Spectroscopy: validation in glioma model and magnetic probe to study cerebrovascular dysregulation in freely-moving Alzheimer’s disease model mice

**DOI:** 10.1101/2021.05.17.444224

**Authors:** Daniel S. Gareau, Michael Bamkole, Matija Snuderl, Cheddhi Thomas, N. Sumru Bayin, Dimitris G. Placantonakis, Julia Zou, Anna Yaroslavsky, Michael P. Dietz, Steven L. Jacques, Sidney Strickland, James G. Krueger, Hyung Jin Ahn

## Abstract

**Significance:** Cerebral vascular reactivity is critical parameters of brain homeostasis in health and disease, but the investigational value of brain oxymetry is diminished by anesthesia and mechanical fixation of the mouse scull.

**Aim:** We needed to reduce the physical restrictivity of hemodynamic spectroscopy to enable Alzheimer’s disease (AD) studies in freely-moving mice.

**Approach:** We combined spectroscopy, spectral analysis software and a magnetic, implantable device to measure vascular reactivity in unanesthetized, freely-moving mice. We measured cerebral blood volume fraction (*CBVF*) and oxygen saturation (S_O2_).

**Results:** We validated that our system could detect delayed cerebrovascular recovery from hypoxia in an orthotopic xenograft glioma model under anesthetized condition and we also found increased *CBVF* and impaired vascular reactivity during hypercapnia in a freely-moving mouse model of AD compared to wild-type littermates.

**Conclusions:** Our optomechanical approach to reproducibly getting light into and out of the brain enabled us to successfully measure *CBVF* and *S*_*O2*_ during hypercapnia in unanesthetized freely-moving mice. We present hardware and software enabling oximetric analysis of metabolic activity, which provides a safe and reliable method for rapid assessment of vascular reactivity in murine disease models as well as *CBVF* and *S*_*O2*_.

## 1. Introduction

### 1.1 Importance of cerebrovascular reactivity

Cerebral perfusion is tightly regulated in response to ischemia (reactive hyperaemia) or the metabolic demand associated with increased local neuronal activation (functional hyperaemia). Cerebrovascular reactivity is therefore a fundamental property of blood vessels and enables rapid adjustment of vascular flow to meet the metabolic needs of brain tissue, and is coordinated by the interaction of neurons, glia, endothelium, smooth muscle cell and pericytes(1). The partial pressure of arterial carbon dioxide (PaCO_2_) is the most potent vascular factor which can induce cerebrovascular reactivity (2). Increased PaCO_2_ (hypercapnia) causes dilation of cerebral arteries and arterioles and leads to an increase in cerebral blood flow (CBF). Alteration of this response impairs the ability to provide sufficient oxygen and nutrients to the brain and leads to disease states (3). Impaired vascular reactivity has been implicated in Alzheimer’s disease (AD)(3, 4). AD patients show decreased cerebral hemoglobin oxygen saturation (*S*_*O2*_) during verbal fluency tasks(5) and a failure of CBF increase during visual stimulus tests (6). Impaired CO_2_ vasoreactivity is associated with cognitive deficits in hypertension patients (7) and blood oxygenation level dependent (BOLD) functional Magnetic Resonance Imaging (fMRI) study showed impaired cerebral vasoreactivity to hypercapnia in patients with AD and amnestic mild cognitive impairment (8).In addition to AD, brain tumors -- particular gliomas -- show high intratumoral vascular heterogeneity with regions of vascular hyperplasia and avascular necrosis interspersed(9). In gliomas, microvascular proliferation in response to tumor hypoxia is closely associated with malignant progression and poor survival(4).

### 1.2 Limitations of Current Technology

Several diagnostic modalities, including fMRI(10), functional Near-Infrared Spectroscopy (fNIRS)(11) and visible light fiberoptic spectroscopy(12), have been used to measure vascular reactivity in animal studies and patients. In animal studies, these techniques require anesthesia in order to minimize movement artifacts, which prevents analysis of cerebral perfusion during behavioral tasks. In addition, anesthesia is known to significantly uncouple neural and vascular responses(13), leading to modified cerebrovascular function. The ability to study changes in cerebral oxygen saturation (*S*_*O2*_) and cerebral blood volume fraction (*CBVF*) without anesthesia could greatly contribute to a more accurate understanding of oxygen needs and cerebrovascular changes in various neurological disorders.

### 1.3 Novel fiberoptic method for measuring S_O2_ and CBVF in freely moving Alzheimer’s Disease mouse model

In order to overcome these limitations, we developed a device and method for measuring *S*_*O2*_ and *CBVF* in freely moving animals using fiberoptic spectroscopy. *S*_*O2*_ depends on the balance of blood flow and oxygen consumption by the tissue. *S*_*O2*_ and *CBVF*, which are both unit-less, together characterize the oxygen content in the tissue. Compared to fMRI, fiberoptic spectroscopy is inexpensive and portable, and offers higher temporal resolution. Compared to fNIRS, visible light spectroscopy penetrates only superficially considering the optical scattering of brain tissue (Fig. 1), offering measurement in a banana-shaped volume between a source fiber which illuminates the brain surface and a radially-displaced detector fiber (Fig. 2). With deeper implantation of the probe fibers, cerebrovascular measurements of deep brain regions, such as the hypothalamus or amygdala, in response to complex behavioral tasks will be possible with our approach. We developed a connection between the mouse brain and the spectrometer consisting of an implanted brain probe (∼1 cm fiber) and a spectrometer connector (∼2 m fiber). The device (Fig. 3a) fiber-optically coupled the brain to the light source and to the spectrometer when magnetically coupled but could be magnetically detached so that the probe could be surgically implanted prior to surgical recovery and subsequent measurement during normal behavior (Fig. 3b,c). After recovery and attachment, spectral measurements were analyzed by least-square fitting to light transport theory (Fig. 3d) to specify the instantaneous *S*_*O2*_ and *CBVF*. These values varied slightly (Fig. 3e) during normal behavior and *CBVF*>0 indicated proper attachment reliably.

**Fig. 1.**
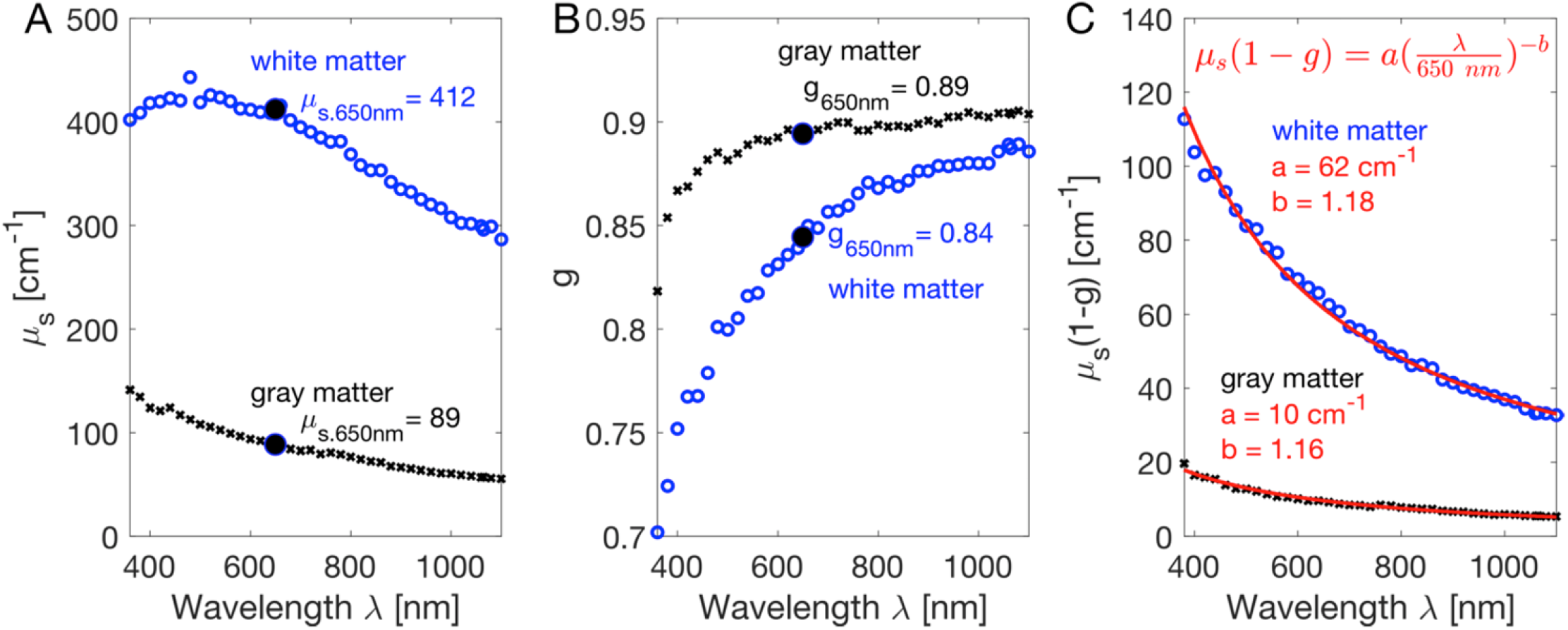
Spectral fit to optical scattering properties. The spectral scattering coefficient (A) and scattering anisotropy (B) from Yaroslavsky et al. 2002 (20) are shown for white matter (blue) and gray matter (black). The reduced scattering coefficient (C) is specified as a fitting function with paramaters a (the μs’ at 650 nm) and b, the scattering power.

**Fig. 2.**
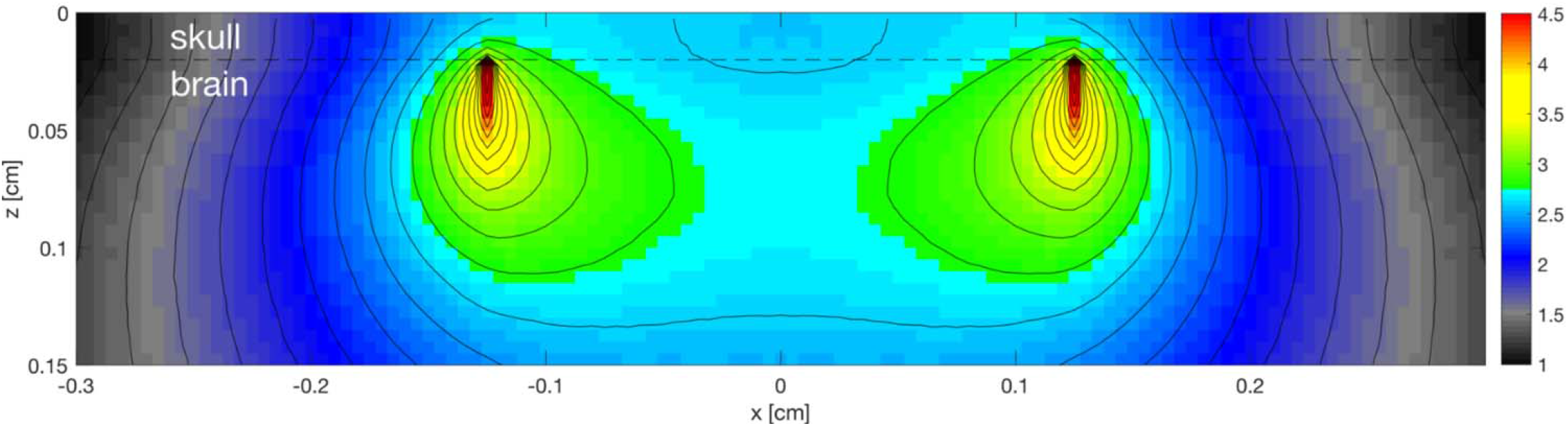
Monte Carlo simulation outputs a map of the of the surgically implanted probe. The light distribution is shown within the murine skull and brain tissue at 650 nm wavelength that is both delivered by the source fiber and collected by the collection fiber. The distribution is the joint probability of delivering to and collecting from each voxel: f_delivered_ x f_collected_, which has units of [cm^-2^]x[cm^-2^] or [cm^-4^], where f denotes fluence [cm^-2^]. The figure plots log_10_(f_delivered_ x f_collected_).

**Fig. 3.**
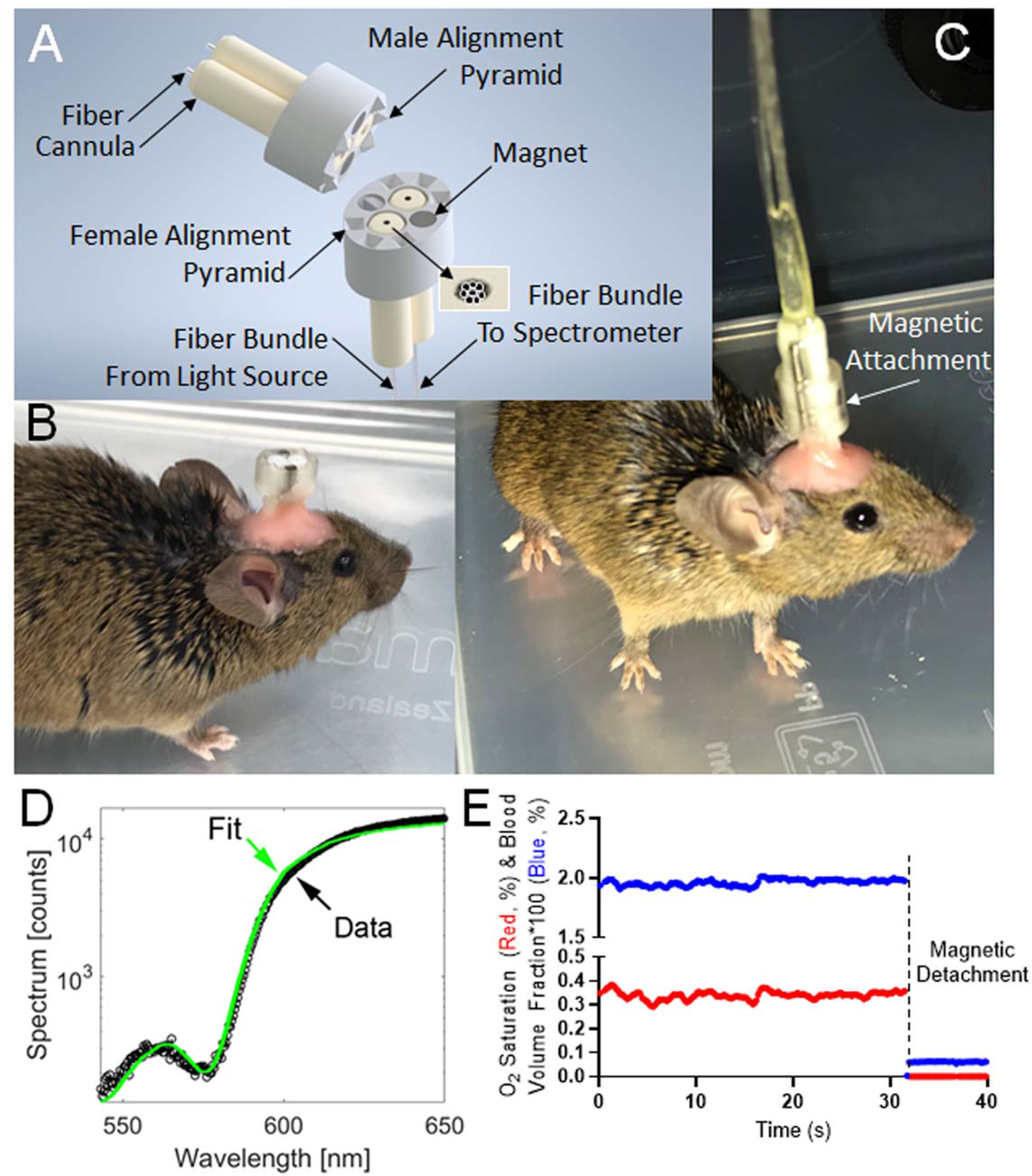
Instrumentation and typical readout for Fiberoptic cerebral oximetry in freely moving mice. (A) 3D model of the magnetic-coupled probe. 6 male/female mating pyramid pairs secure the detachable, implantable brain probe into a linking fiberoptic with mating female pyramid components to enable fiberoptic alignment. (B) surgically-implanted brain probe. (C) brain probe connected to a spectrometer. (D) spectral fitting in <0.25 seconds yielding instantaneous measurements of the percentages of *CBVF* and *S*_*O2*_ in the 540nm to 650nm range chosen to contain 3 isosbestic points of hemoglobin spectral absorption alongside an anchoring wavelength (650nm) of differential absorption. (E) Typical time-course measurement over resting behavior and decoupling event corresponding to magnetic probe detachment. The graph shows cerebral blood volume fraction (blue) times 100 such that a y-axis value of 1 means 1% blood volume fraction and the oxygen saturation fraction (red), which is the fraction of hemoglobin molecules bound to oxygen within the cerebral probed volume.

## 2. Materials and Methods

### 2.1 System and computational method of fiberoptic measurement

A white light source (HL-2000-HP, Ocean Optics, Dunedin FL) illuminated the cerebral cortex through an illumination fiber bundle of 8 fibers (FG105UCA, Thorlabs, Newton, NJ) in 7-around-1 mated to a solid core fiber in the tip (FT400UMT, Thorlabs, Newton, NJ), while an identical detection fiber tip and bundle directed the measured diffuse reflectance to a spectrometer (HDX, Ocean Optics, Dunedin FL). Our system was controlled through Matlab (Mathworks, Natick, Massachusetts) software by a laptop computer running Windows 10. Spectra were fit for real-time readout of *CBVF* and *S*_*O2*_ on the perioperative laptop during experimentation. The graphic user interface is provided in the supplementary materials. The sampling rate of the spectroscopic measurement (generally ∼1Hz) is set determined by the spectrometer integration time, which is automatically set over iterative measurements during spectroscopy to achieve a signal that is in the high, but not saturated region of between 70% and 90% of the spectrometers’ specified saturation level.

### 2.2 Spectroscopy

Calibration spectra *C*(λ) were acquired using measurements of a 99% reflectance standard (AS-011XX-X60, Labsphere, North Sutton, NH) while holding the fiber probe 3 cm from the standard. The dark noise of the spectrometer was also measured (*dark*) with the illumination lamp off. Calibrated spectral measurements *M*(λ) of tissue were calculated from the raw tissue spectral measurements *R*(λ) and the calibration measurements *C*(λ) according to **Equation 1**. Measured spectra depended on the spectra of the light source and the detector sensitivity but these factors were cancelled by the normalization:

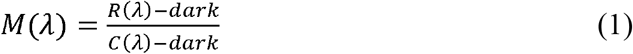

A theoretically predicted spectral measurement *M*_*p*_(*λ*) was created by multiplying the predicted reflectance, *R*_*p*_(*λ*), by a scaling factor K: *M*_*p*_(*λ*) = *KR*_*p*_(*λ*) as in equation 13, below. Least squares minimization between *M*_*p*_(*λ*) and *M*(*λ*) specified the endpoint metrics *S*_*O2*_ and *CBVF*. Equations 3-13 implement diffusion theory (14-17) to calculate R_p_(*λ*) as adopted from our previous esophagus studies (12, 18).

Equations 3-12 are implemented in the simulated diffuse reflectance spectrum in the supplementary Matlab spectroscopy code (GUI.m) and the key diffusion parameters are defined in the referenced literature, such as the internal reflection coefficient *A* for light totally internally reflecting within the media at the media boundary, the equivalent point source depth *z*_*0*_(λ), diffusion constant *D*(*λ*) and the effective attenuation coefficient *μ*_*eff*_(*λ*). Diffusion theory was considered valid since the 2.5 mm source-detector separation on our probe was large compared to the transport scattering mean free path (*MFP’* = 0.39mm), calculated using scattering spectra (Supplementary Figure 2A and 2B) and for the least scattering wavelength in our measurement (650nm). The reduced mean free path (*MFP’*) is effectively the probing depth of optical penetration.

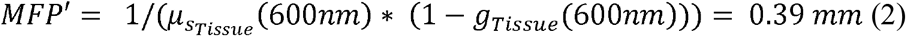

where “Tissue”denotes the approximate mixed brain tissue type to be 1/3 white matter and 2/3 gray matter as in Equation 14 and Equation 15, below.

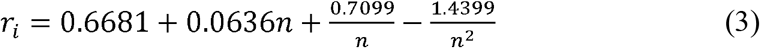

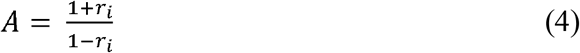

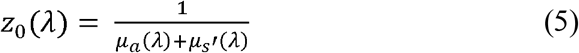

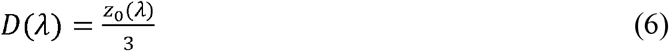

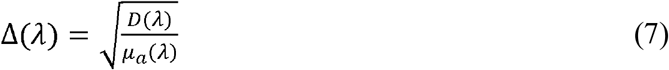

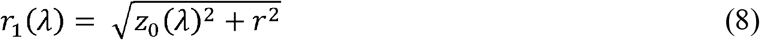

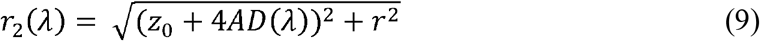

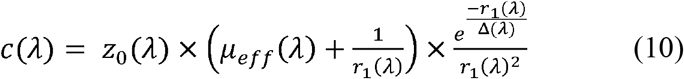

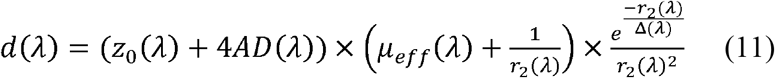

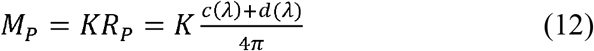

The fiber separation (distance between the source fiber tip and the detector fiber tip) was *r* ≈ 0.25 cm, and the refractive index mismatch between glass (n_glass_ = 1.52) and tissue (n_tissue_ = 1.35) was *n* = n_glass_/n_tissue_ = 1.09. *μ*_*a*_(*λ*) and *μ*_*s*_’(*λ*) were the absorption and reduced scattering coefficients of the tissue, respectively. The spectral absorption coefficient, *μ*_*a*_(*λ*), was calculated as a weighted sum of the absorption spectra of oxygenated whole blood (*μ*_*a_oxy*_(*λ*)), deoxygenated whole blood (*μ*_*a_deoxy*_(*λ*)), and water (*μ*_*a_water*_(*λ*)), according to Equation 13:

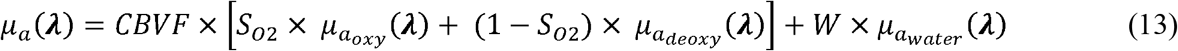

### 2.3 Calculation of Fitting Parameters to Quantify SO2 and CBVF

Spectra were analyzed in the range 540 to 650 nm by least-squares fitting. The rationale for this wavelength range is that the optical penetration is restricted to the superficial brain layers including the cortex (i.e. not too long) and that the range includes multiple isobestic points of the hemoglobin absorptioin curve so it is rich in information content regarding hemoglobin saturation.

The difference between *M* (Eq. 1) and *M*_*p*_ = KR_p_ (see Eq. 12) was minimized by adjusting *M*_*p*_ using a multidimensional unconstrained nonlinear minimization method (Nelder-Mead), the fminsearch() in Matlab. Fitting adjusted 3 variables: *CBVF, S*_*O2*_ *and K* where *CBVF* = 1 specifies 150 g/L hemoglobin. *W*, the fractional tissue water content, was fixed at an assumed value of 0.78, which has been reported for mouse cortex (19). Absorption due to water was minimal. For example, at 650 nm, low *CBVF* = 0.01 and *S*_*O2*_ = 0.5, the absorption coefficient in the tissue due to the presence of blood is *μ*_*a*_blood_ = *CBVF*×*S*_*O2*_×μ_*a*_oxy_ + *CBVF*×(1-*S*_*O2*_)×*μ*_*a*_deoxy_ = 0.1103 cm^-1^. *μ*_*a_water*_ = 0.0025 cm^−1^, which is 44-fold lower than *μ*_*a*_blood_.

The absorption of the tissue due to blood was specified by the fitting described above while the scattering optical properties (scattering coefficient *μ*_*s*_(*λ*) [cm^-1^] and scattering anisotropy *g*(*λ*) [-]), which were constant across all experiments, were spectral functions (*λ*) fit to the previous measurements of fresh human brain tissue (20). Thus *μ*_*s*_(*λ*) and *g*(*λ*) were used as interpolated functions of wavelength (**Equations 14-15)** that, in combination yielded the reduced scattering coefficient, *μ*_*s*_’(*λ*) = *μ*_*s*_(*λ*)×(1-g(*λ*)) used in **Equation 5**. These optical properties agree reasonably with the literature (20, 21), such as our previous data (20) that can be fit for the reduced scattering coefficients of gray and white matter as shown in Fig. 1C.

### 2.4 Brain optical scattering properties

Fig. 1 shows the brain scattering properties specified by the spectral data from Yaroslavsky et al. 2002 (20). The Fig. 1a and Fig. 1b show the spectral behavior of the scattering coefficient (*μ*_*s*_) and the anisotropy of scattering (g). The combination of *μ*_*s*_ and g yields the reduced scattering coefficient (*μ*_*s*_’ = *μ*_*s*_(1-g)) shown in Fig. 1c for white and gray matter. Scattering properties of whole tissue were set as a weighted-sum based on a 1:2 ratio of white to grey matter, as reported (22) for mouse cortex.

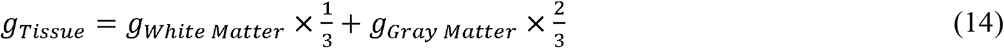

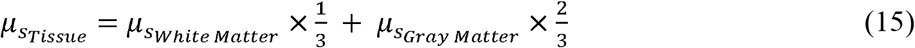

The *μ*_*s*_’(λ) spectrum was assumed when using least-square fitting of the data to specify *K, CBVF* and S_O2_. After spectral fitting, the values for *CBVF* and *S*_*O2*_ were output as a function of time (e.g. **Fig. 1e**) over time periods where spectroscopic measurements were made on mice under various experimental conditions. These optical properties were also used to model the photon transport in the mouse brain.

### 2.5 Fabrication of fiberoptic probe with surgically implantable, magnetically-coupled tip

The connection between the mouse brain and the spectrometer consisted of an implanted brain probe (∼1 cm fiber) and a spectrometer connector (∼2 m fiber). The connector (Fig. 3a) fiber-optically coupled to the illumination source and the spectrometer and opto-magnetically coupled to the brain probe, which fiber-optically coupled the brain.

Plastic connectors were 3D-printed using a 3D Systems Projet printer and Visijet M3 Crystal material. Mating surfaces were 3D-printed facing upward (opposite the print bed) to minimize warp and maximize surface flatness. The plastic connector design was optimized to align and hold (press fit) two ceramic ferrule cannulas (CF440-10, Thor Labs, Newton NJ) and two cylinder magnets (D12-N52, K&J Magnetics, Plumsteadville PA).

During assembly, the connector’s flat surface was pressed against glass, and the ferrules and magnets were inserted and also pressed against the glass for alignment (friction fit). The male side connector was fabricated using a similar process of press fitting the cannulae and magnets, but the assembly was pressed against an alignment tool (see Supplemental Materials), instead of the flat glass plate. The connector was then mated to its probe with magnetic attachment, and pre-polished fibers (FT400UMT, Thor Labs, Newton NJ) were inserted into the ferrules and affixed using UV-cure adhesive (#68, Norland Products, Cranbury NJ). After dry, the protruding fibers were clipped and polished to a length of 0.5 mm protruding from the cannulas using a custom polishing tool set that held the probe such that the fibers were normally incident on standard fiber polishing film.

In order to maintain alignment, so that the fibers in the brain probe and spectrometer connector mated precisely, the brain probe surface was printed with 6 small 4-sided pyramids while the connector surface was printed with complementary mating holes. This design configuration enabled secure mating of the two connecting parts during measurements. Finished fiber probes were visually inspected under 20X stereomicroscopy for polish level and analyzed through a series of signal and noise tests using the spectroscopy probe in a dark room and the light source targeted on human skin.

The quantitative performance parameter for magnetic coupling was the amount of illumination white light that leaked directly from the source fiber bundle to the detector fiber bundle. This amount defined the noise and was measured with the surgically-implantable tip magnetically coupled and the illumination light going from it toward the Vantablack target in a dark room. The quantitative spectrometer readout, in counts, was read out (***R1***) at the 650nm wavelength. A second count readout (***R2***) was measured with the probe contacting human skin. The ventral skin surface is less pigmented and the easily-accessible, finger pad has little unwanted influence of melanin. The signal to noise ratio was then [***R1*** – ***R2***] / ***R1*** and manufacturing quality control threshold set as a 60-1 signal-to-noise ratio so that, as a first order approximation, an extra 1/60^th^ of an optical measurement of brain consisting of leaked white light would erroneously decrease the absolute *CBVF* no more than 100%/60 = 1.6%.

### 2.6 Analysis of cerebrovascular reactivity of freely moving mice

5-month-old C57BL/6 mice (The Jackson Laboratory) were used to measure basal cerebrovascular dynamic properties of the cortex while freely moving. The AD model mice studied were 13-month-old Tg6799, also called 5xFAD (Jackson Laboratory) double transgenic mice overexpressing both human amyloid precursor protein (APP) gene with KM670/671/NL, V717I, and I716V mutations and human presenilin 1 (PSEN1) harboring M146L and L286V mutations under the Thy1 promoter (23). Wildtype (WT) littermates were used as controls. Only male mice were used in experiments. All mice were maintained on a 12-hour light/dark cycle and given ad-lib access to irradiated mouse chow and water for the duration of the experiment. All experiments were done according to policies on the care and use of laboratory animals of the Ethical Guidelines for Treatment of Laboratory Animals of the NIH. Relevant protocols were approved by the Rockefeller and Rutgers Institutional Animal Care and Use Committee (IACUC).

After anesthesia with isoflurane mixed with pure oxygen (3% isoflurane and 1 L/min oxygen for induction and 1.5% isoflurane and 1 L/min oxygen for maintenance), mice were mounted on a stereotactic frame. After the removal of hair over the scalp, the scalp was cleaned with sterile alcohol and Betadine three times. We performed a minimal-sized midline scalp incision (∼5 mm). We then drilled two small holes in the skull using a 0.7 mm diameter stainless steel micro drill burr (Fine Science Tools). The coordinates of the two holes were Bregma= -1 mm, lateral= 1.5 mm and Bregma=-3.5 mm, lateral=1.5 mm. Brain probes were fitted to the holes and immobilized using dental cement. The skin incision was sutured and animals were monitored throughout the recovery period. A week after surgery, mice were placed into a chamber (22.8 cm x 20.3 cm x 15 cm) and we measured cerebral perfusion in the freely moving state. For the hypercapnia study, we injected 10%CO_2_/10% O_2_/80%N_2_ mixture into the induction chamber and measured changes in *CBVF* and *S*_*O2*_ during and after hypercapnia.

### 2.7 Spectroscopy validation by analysis of cerebrovascular reactivity in mouse model of glioma

A separate group of mice were used under anesthesia for the glioblastoma model at the New York University. These mice were housed within NYU School of Medicine’s Animal Facilities. Brain tumor implantations were performed according to protocols approved by NYU’s IACUC, as previously described (24). Briefly, 6-8 week-old NOD.SCID (NOD.CB17-Prkdcscid/J, 001303, Jackson Laboratory, Bar Harbor, ME) male mice were anesthetized with intraperitoneal injection of ketamine/xylazine (100 mg/kg and 10 mg/kg, respectively). They were then mounted on a stereotactic frame (Harvard Apparatus). A midline skin incision was made. A high-speed drill was used to drill a small hole in the calvaria 2 mm off the midline and 2 mm anterior to coronal suture. Five μl of a suspension of U87 cells (100,000 cells/μl) were injected through a Hamilton syringe (1 μl/min, Harvard apparatus, needle pump) into the left frontal lobe through the drilled hole The injection needle was left in place for an additional 5 min after the injection was completed to prevent backflow. The skin incision was sutured and animals were monitored throughout the recovery period. U87-MG cells for production of intracranial xenografts were cultured in Dulbecco’s Minimal Essential Media (DMEM, Life Technologies) supplemented with 10% FBS and non-essential amino acids. Previous studies with U87-MG cells in our laboratory showed large tumor formation 2 weeks after injection (24). Two weeks after the injection of cells, terminally ill animals were prepared for fiberoptic measurement. After anesthesia and fitting into the stereotactic frame as described above, we drilled an additional hole in the contralateral frontal calvaria, using a high-speed drill. The diameter of the 2 holes matched the caliber of the probe. After cleaning the area with ethanol wipes, the probes were fitted to each window and immobilized using dental cement. Animals were removed from the stereotactic frame for fiberoptic measurements. We first measured *CBVF* and *S*_*O2*_ at the resting state for 30 sec under ketamine/xylazine under anesthesia. To analyze the cerebrovascular response during hypoxia, we blocked the nostrils for 30 sec with a wet cotton swap, and measured *CBVF* and *S*_*O2*_ during hypoxia. After nostril blockage release, we continued *CBVF* and *S*_*O2*_ recordings during reperfusion. This procedure was repeated once, then animals were euthanized following the approved protocol. The skull was opened and the presence of the glioma in situ under the probe was confirmed by a neuropathologist.

## 3. Results

### 3.1 Analysis of cerebrovascular reactivity in freely moving C57BL/6 and AD mice

Fiber spectroscopic measurement of *CBVF* and *S*_*O2*_ in mixed arterial and venous blood the cortex of freely moving 5-month old C57BL/6 mice yielded values of ∼1.9% and ∼ 35% respectively (Fig. 4a). To investigate the change in *CBVF* and *S*_*O2*_ of the cortex during hypercapnia in the mouse cortex, we placed mice in an insulated chamber and increased CO_2_ concentration up to 10%. When hypercapnia was induced after 60 seconds of baseline measurement by increasing CO_2_ concentration, we found an increase in *CBVF*, which is a typical hyperaemic response to increased CO_2_ through vasodilation. *S*_*O2*_ decreased slowly but normalized rapidly after stopping CO_2_ injection and releasing of CO_2_ at 300 sec (Fig. 4b), while *CBVF* increased rapidly before returning to a level that was less than the baseline measurement.

**Fig. 4:**
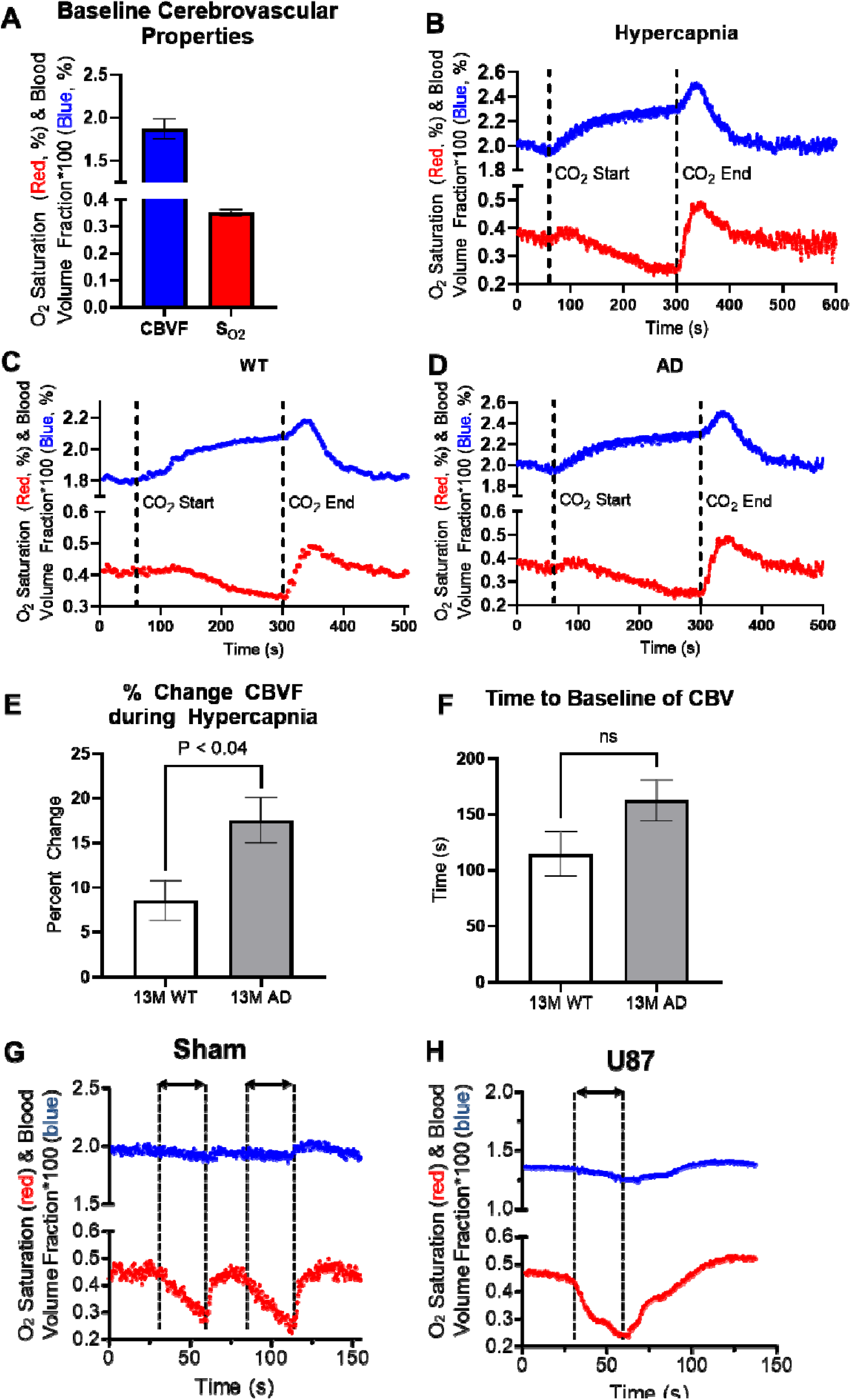
In vivo spectroscopy results. (A), Basal cerebrovascular properties of the cortex of a freely moving, 5 month-old mouse. (B) Cerebrovascular responses in 5 month-old freely moving mice were obtained using fiberoptic spectroscopy during hypercapnia. Data are representative of three independent experiments. (C) & (D), Representation for cortical vascular reactivity of Tg6799 13 month-old mice (C) and WT littermate (D) during hypercapnia were analyzed using fiberoptic spectroscopy. Hypercapnia was induced at 70 sec by increasing CO_2_ concentration up to 10 % and was kept until 300 sec. Data are representative of four independent experiments. (E), The increase of cerebral blood volume fraction (*CBVF*) during hypercapnia was significantly higher in Tg6799 mice compared to WT littermate (p<0.04, n = 4 AD, 4 WT). (F) There is a tendency of delayed time to the baseline of CBVF after hypercapnia in Tg6799 mice, but it is not statistically significant (n = 4 AD, 4 WT). *(G)*, Hypoxic conditions were held from 30 – 60 sec and from 80 – 110 sec in 2 month-old mouse. The graph shows cerebral blood volume fraction (blue) and the oxygen saturation fraction (red). (H), Delayed Cerebrovascular recovery from hypoxia in a NOD.SCID 2 month-old mouse with intracranial U87 GBM xenograft was delayed compared to a sham mouse. Bar graphs represent mean ± s.e.m. of ≥3 independent experiments, and was statistically analyzed by performing a two-tailed unpaired t-test.

To investigate the cerebrovascular properties of an AD (Tg6799) (23) mouse, we measured the baseline of *CBVF* and *S*_*O2*_ of the cortex of freely moving 13 month-old Tg6799 mice and wild-type (WT) littermates and also analyzed the difference of vascular reactivity between Tg6799 mice and WT littermates during and after hypercapnia using fiber optic spectroscopy (Fig. 4c,d). While there is was no difference in baseline of *CBVF* and *S*_*O2*_ between the two groups (Fig. S1), the increase of CBV of the cortex of AD mice during hypercapnia was significantly higher than that of WT littermates (Fig. 4e). In addition, although not statistically significant, the time to baseline of *CBVF* of AD mice was also delayed in comparison to WT littermates (Fig. 4f).

### 3.2 Validation of impaired cerebrovascular reactivity measurement using glioma mouse model

To validate our method and demonstrate broad utility, we analyzed cerebrovascular response to hypoxia in an orthotopic xenograft mouse model generated with human U87 glioblastoma (GBM) cells. Since patients with brain tumors show impaired cerebral vasoreactivity (25-27) and hyperproliferation of immature blood vessels lacking smooth muscle cells limiting perfusion in malignant glioma (4), an orthotopic xenograft glioma mouse model is a good platform to validate our new system. Two weeks after orthotopically injected with human U87 GBM cells, the fiberoptic probes were surgical implanted on the skull of glioma and control mice (sham injected) and we induced hypoxia for 30 seconds and measured the *S*_*O2*_ and *CBVF* change directly from brain parenchyma under ketamine/xylazine anesthesia. In sham mice, during hypoxia for 30 sec, cerebral *S*_*O2*_ decreased but normalized within 30 seconds after hypoxic release (Fig. 4g). We observed the same pattern of hypoxia and rapid normalization at each cycle of obstruction/release as expected for a normal cerebrovasculature, but the glioma mouse had a recovery of more than 30 seconds to recover their normal *S*_*O2*_ (Fig. 4h). This result suggests that our system could detect cerebrovascular alteration and impaired vascular reactivity in glioma mouse model.

## 4. Discussion

Fiberoptic spectroscopy can assess the *S*_*O2*_ and the *CBVF* in the brains of freely moving mice when implemented with a magnetically coupling tip. The present study showed that our fiberoptic method detected changes in freely moving mice and measured cerebrovascular impairment in a mouse model of AD. *S*_*O2*_ in cerebral blood of control mice was 30 - 60%, which is comparable to previous studies in the cortex of anesthetized rodents (28, 29). *CBVF* of 6-8 week old NOD.SCID mice was ∼2.0% (Fig. 4g), *CBVF* of 5 month-old C57BL/6 mice was 1.9% (Fig. 4a,b) and *CBVF* of 13 month-old Tg67999 (C57BL/6 background) was ∼ 1.7% (Fig. S1). This range from 1.7% to 2% in our results is slightly less than the range reported in the literature: 2.1% reported by Synchrotron Radiation Quantitative Computed Tomography in the parietal cortex (30), 2.03-3.14% reported by MR in the cortex (31-33), 2.0-2.4% reported by multiphoton laser scanning microscopy and in capillary-rich cerebral cortex (34). One possible reason is that we measured *CBVF* of 5-month old and 13 month-old mice, instead of young rodents (4-8 weeks old), which other studies mainly used (31-34). Although we cannot directly compare because of different strains, the young NOD.SCID mice (8-10 weeks) in the GBM experiment (Fig. 2g) showed higher *CBVF* values than 5-month old and 13-month-old mice (Fig. 2a,b). Another possible reason is that large blood vessels on the surface of brain such as meningeal vessels were excluded from our assessment because our fibers were implanted ∼ 0.3 mm below the skull.

Tg6799 mice starts to develop amyloid plaque deposits by 2-month-old age (23) and shows substantial amount of amyloid plaque deposits and cerebral amyloid angiopathy in their cortex (35). In addition, recent dynamic susceptibility contrast-enhanced (DSC) MRI analysis showed increased vasodilation in 12-month-old Tg6799 mice compared to littermate control, in response to contrast injection (36). This result is quite similar to current study showing increase vascular reactivity in Tg6799 mice. Although cerebral hypoperfusion is typically observed in advanced AD patients(37, 38), an increase of cerebral perfusion was also seen in some cases of early AD patients (39-41). Integrating several studies of cerebral blood flow (CBF) in AD patients suggests that there is cerebral hyperperfusion in early disease followed by hypoperfusion later in the disease (42, 43). In AD patients, as disease progresses, severe neuronal cell death and brain atrophy have been widely reported (44, 45). However, our AD mouse model (Tg6799) showed very minor neuronal cell death in limited brain areas and no brain atrophy (46). Therefore, Tg6799 may represent the early stage of disease and hyperperfusion of this mouse model could be relevant to early AD patients.

## 5. Conclusions

This fiberoptic method may improve understanding the mechanisms of impaired vascular reactivity in neurologic disorders and brain tumors. Our vasculature and hemodynamic measurements raise the question as to whether the observed phenomena are consequences of the pathology or more fundamental changes that are required for disease progression. We hope to explore these temporal measurements in future work, potentially combining *CBVF* and *S*_*O2*_ with time-differential mathematics to access perfusion and oxygen consumption parameters. Understanding the extent of impaired vasoreactivity in brain pathologies may someday enable novel diagnostic and prognostics without need for ex-vivo studies. Though reliable relative S_O2_ and *CBVF* values over longitudinal measurements have been shown here, a current limitation is the lack of validation of absolute estimates of these parameters across animal models, and results thus far may be subject to estimation error.

Mechanically, our device allows natural animal movement because the flexibility of our fiber bundle and a video of exemplary animal free movement is provided as a supplementary material. When animal movement becomes so severe that continued connectivity of the device would become dangerous to the animal, our magnetic detachment clean brake-away protects the animal and therefore measurement integrity of future measurements. Figure 1E shows the distinction between resting fit measurements and the fitting result when the magnetic attachment (which assures fit quality) has been broken. In our experience, when the magnetic coupling is attached, accuracy can be assured in terms of the quality of fit as evidenced (but not explicitly shown in this paper) across our entire study, and it is clear when that condition is broken, as evidenced in Figure 1E. Comparatively, other designs do not lock into the biometric (spectral fit quality) with the same degree of detachment simplicity, and this leads to not only the animal being less-free and potentially being injured by the apparatus if it should not detach easily but also leads to the application of torque to the head corrupting the data when such a design without magnetic coupling/detachment is in a state of mechanical stress or pressure. Quantitative evidence for this advantage is expected in ongoing comparative studies.

## Disclosure/ Acknowledgments

The Authors have no relevant financial interests in this article and no potential conflicts of interest to disclose. This work was supported in part by the National Institute of Health NS104386 and UL1 TR000043-09 (Ahn) and in part by the Clinical and Translational Science Award (CTSA) program. Grant Number: UL1 TR001866. Placantonakis was supported by NINDS R01 NS102665 and NYSTEM IIRP DOH01-STEM5-2016-00221/C32595GG.

## Supplementary Materials

The 3D models to print our device, the commercial component part list, and the operating system software to replicate our work are available in the public repository at Rutgers University Libraries (https://scholarship.libraries.rutgers.edu/esploro/outputs/technicalDocumentation/Code-data-and-materials-for-Fiberoptic/991031663984304646?institution=01RUT_INST). The software supports the demonstration of the fitting of an included sample spectrum as well as the acquisition of a time-series of spectral measurements that it will fit in real-time.

## Notes

https://rockefeller.box.com/s/r9x5qmfgvpo7fsq325h2pwtfxqldl07r

### Competing Interest Statement

The authors have declared no competing interest.

### Summary of Updates

We identified an error our MatLab code for the spectroscopic measurement algorithm: the refractive index was incorrect, and this seems to have affected the absolute value of the CBVF measurements reported in the original manuscript. We corrected this error in our software implementation and reanalyzed the data collected in the open arena and hypercapnia measurements of 13-month-old Tg6799 mice and wildtype littermates. The average CBVF reported among the animals was found to be around 1.7%.

