## Supplementary Materials for "Fiberoptic Hemodynamic Spectroscopy: validation in glioma model and magnetic probe to study cerebrovascular dysregulation in freely-moving Alzheimer’s disease model mice"

### Supplementary Figure

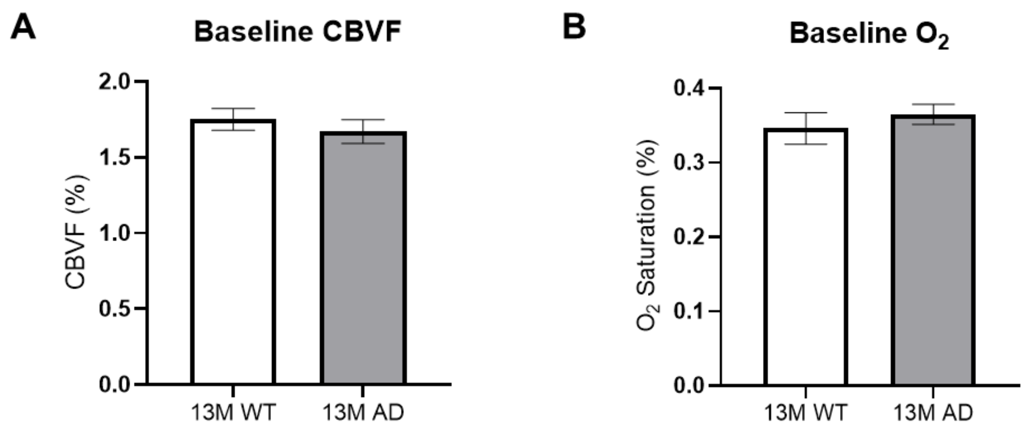


**Figure S1. Basal cerebrovascular properties of freely moving 13 month-old AD mice and WT littermates.** 13 month-old Tg6799 mice and WT littermates showed similar levels of basal oxygen saturation (a) and cerebral blood voume fraction (b) in the cortex. ($p > 0.05$, n = 4 AD, 4 WT)

### Detailed Instructions For Fiberoptic Fabrication

**Materials**

Probe fiber (Ø400 µm Core Multimode Optical Fiber): FT400UMT (Thor Labs)

Connector fiber (Ø105 µm Core Glass Clad Silica Multimode Optical Fiber): FG105UCA (Thor Labs)

Ceramic ferrules (Ø2.5 mm, 10.5 mm Long Ceramic Ferrule for MM Fiber, Ø440 µm Bore): CF440-10 (Thor labs)

Magnets (1/16"x1/8" cylindrical neodymium magnets): D12-N52 (K&J Magnetics)

Norland Optical Adhesive 68: NOA68 (Norland Products)

Glue for SMA connectors: 5 Minute Epoxy (Devcon Home)

Probe plastic frame: Attached STL and Inventor CAD files (printed on 3D Systems Projet with M3 Crystal)

Connector plastic frame: Attached STL and Inventor CAD files (printed on 3D Systems Projet with M3 Crystal)

**Tools**

Fiber optic stripper tool for 105 micron fiber: Micro-Strip .008 MS1-08S-13-FS (Micro Electronics)

Fiber optic stripper tool for 400 micron fiber: Micro-Strip .021 MS1-21S-40-FS (Micro Electronics)

Diamond polishing films (for fibers, ferrules and SMA connector tips): LF30D, LF6D, LF3D, LF1D, LFCF (Thor Labs)

SMA Connector Polishing Disc: D50-SMA (Thor Labs)

Fiber/Ferrule Polishing Disc: D50-F (Thor Labs)

Glass polishing plate with silicone pad: CTG913 , NRS913A (Thor Labs)

UV Curing light: AccuCure Spot ULM-3B-365 (Digital Light Lab UV-LED Curing Systems)

Dark standard: VantaBlack VBS-3555 (Surrey Nanosystems)

SMA-905 Connectors (Thor Labs)

Fiber Optic Kevlar Shears: Ripley Miller 86 1/2SF (Specialized Products)

25 blade metric feeler gauge: W80527 (Performance Tool)

Flat metal mounting base: BA1 (Thor Labs)

Translation Stage with Standard Micrometer: PA1 (Thor Labs)

Probe Assembly polishing tool: Attached CAD files (printed on Carbon 3D with CarbonResin PR25)

**Equipment**

Spectrometer 1: HR2000+ with 200-1100nm Grating HC1, ILX-511B Detector DET2B-200-1100 & 25µm Slit SLIT-25 (Ocean Optics)

Spectrometer 2: Ocean-HDX-VIS-NIR with 50µm Slit INTSMA-050 (Ocean Optics)

Halogen Light Source 1: HL-2000-HP-FHSA (Ocean Optics)

Halogen Light Source 2: HL-2000-HP (Ocean Optics)

Stereo microscope: Zeiss Stemi 2000

**Software**

PC with Windows 10 (Microsoft), MATLAB and Instrument Control Toolbox (Mathworks), Spectrasuite (or Oceanview) and OmniDriver (Ocean Optics)

**Fabrication Instructions: Probe Fiber Sections**

1. Cut sheet of cardboard or paper to length (14mm) to use as a guide for cutting fiber sections
2. Hold length guide made in previous step against Probe Fiber (FTUMT400) and cut many fiber sections with Fiber Optic Shears (Ripley). Store cut sections in a container labeled "not polished."
3. Place green polishing film (LF30D) on silicone pad (CTG913 , NRS913A) and place several drops of distilled water with a transfer pipet to wet the whole film
4. Use fingers to spread distilled water across and flatten polishing film (LF30D), rub from the center outward to minimize air bubbles between silicone pad and film. If too many bubbles remain, lift film or part of film and reapply until bubbles are minimized
5. Insert ferrule (CF440-10) into polishing disc (D50-F) and insert not polished fiber section into ferrule
6. Fold a sheet of laminated paper (as found in product packaging eg alcohol wipe packets) a few times to place between index finger and raw fiber while polishing (this folded sheet should protect finger from being cut by fiber while polishing)
7. Apply slight downward pressure to fiber with index fingertip (with a force of about 40-90 grams) while moving the polishing disc in figure-8 motions along the polishing film. Start with about 5 figure 8 motions, then follow with about 5 more figure 8 motions in the opposite direction
8. Check the level of polish using stereoscope at 30-50x. If the fiber surface is not flat, return to the green polishing film(LF30D). If the fiber is flat, then proceed to the red polishing film (LF6D). Repeat polishing and check fiber under stereoscope for each film (LF3D,LF1D and LFCF). After each film is used, clean it with a dry task wipe or paper towel and reapply distilled water before each use. Before checking the fiber for final polish, clean the fiber with a wet (or alcohol wipe) then a dry task wipe. Store polished sections in a container labeled "fine polished."

**Fabrication Instructions: Probe Assembly**

1. 3D print (3D Systems Projet M3 Crystal) the probe assembly male and female connectors as well as the probe polishing tools (3D Systems Projet Crystal or Carbon 3D). Orient the connectors' mating surfaces upward on the print bed so that no scaffold support wax is used near the parts' connecting features. Clean the connectors in ultrasonic cleaner for 20-30 minutes to remove all wax.
2. Lay a ¼” hex tool on flat work surface then place the male connector on upward face of hex tool, with the male features facing down toward the table. Press ferrule (CF440-10) into the connector until it clicks against the hex tool (ferrule should be aligned flush with connector mating surface, verify flush alignment under stereo microscope). Clean the ferrules using an alcohol wipe then compressed air.
3. Place magnets (D12-N52) on connector to check orientation, then lightly insert magnets into probe assembly.
4. With the connector female end in a clamp pointing upward, place the male probe assembly on top of the connector end and press the probe's magnets all the way in.
5. Press lightly on the male probe assembly to make sure it is fully seated against the female connector end. Check two finely polished fibers to make sure they are fully polished and cleaned, then insert the fibers into the ferrules. (The connector can be illuminated during insertion to visually verify proper insertion).

6. Lift each fiber about 3-4mm. Use a cut section of clad fiber to apply optical adhesive (about 2-3 microliters) around the fiber, then reinsert each fiber fully into its ferrule. Spray away excess adhesive with compressed air pointing perpendicular to protruding fiber tip. Wipe off any extra remaining adhesive with a task wipe. Using a cut section of clad fiber, apply optical adhesive to the top perimeter of each magnet/magnet hole sidewall. Cure the probe adhesive while probe remains attached to the connector in the clamp using UV light (expose for 45-90 seconds, use UV safety eye protection).
7. Insert the probe into the probe polishing tool (length 1 +0.85mm) and press on the probe with index finger and a small piece of task wipe, holding the probe in the polishing tool while polishing (in figure-8 motions) on the green film (LF30D). The film should be wet with distilled water. The sound will change as the fiber is polished to length (the vibration of the polishing assembly will also feel different).
8. Clean the probe tip with a task wipe and wipe the polishing film dry with a paper towel. Insert the probe into the next polishing tool (length +0.65mm) and polish on the red film (LF6D) to length.
9. After each film, wipe the tip and film, and check the tip under stereoscope. Use the third probe polishing tool (length +0.5mm) while polishing 5 figure-8 motions in each direction on each remaining film (LF3D, LF1D & LFCF).
10. Measure the final fiber protruding tip length visually with feeler gauge to the nearest 0.1mm. Record the tip length along with signal and noise for each probe. Wrap the polished probe in a task wipe and store in a labeled vial.

**Figure S1: Polishing Tool**

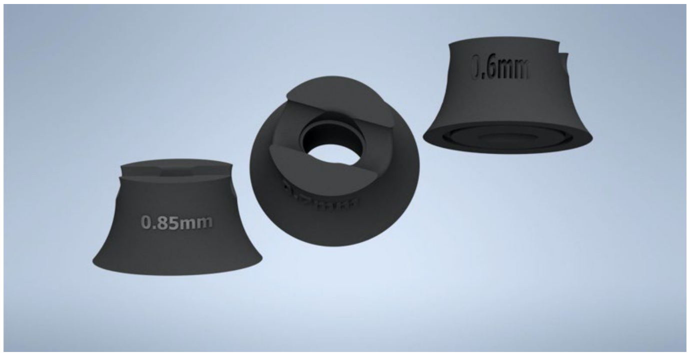


**Fabrication Instructions: Connector**

1. Cut 16, equal lengths of fiber (FG105UCA), with the length being long enough to span the distance from the spectrometer and light source to the animal enclosure
2. Strip 8 ends about 12-13mm long using Micro-Strip .008 (MS1-08S-13-FS). Insert stripped ends into ferrule (CF440-10). Verify all fiber tips insert past the length of ferrule. Pull bundle out about 3-4mm, then apply optical adhesive and reinsert. Apply additional optical adhesive to protruding fiber tip end of cannula. Cure using UV light and eye protection.
3. Polish ferrule end using D50-F, set depth in D50-F using jig and tightening set screw. Continue to re-set depth in jig while polishing through the films (LF30D-LFCF). Repeat process from step 2 for second bundle of 8 fibers.
4. Strip the 8 ends of a fiber bundle 15mm while keeping the total length of all 8 fibers approximately equal. Place SMA connector in a clamp. Insert stripped fibers through SMA connector strain relief shroud and sleeve then fix fibers with epoxy into SMA connector. Press SMA strain relief shroud onto connector and let cure. This 8 fiber bundle can be labelled as the "illumination side." Polish using

D50-SMA and polishing films (LF30D-LFCF) with distilled water as in previous polishing steps above.

5. Strip the remaining 8 fiber bundle ends about 13-14mm while keeping the total length of all 8 fibers approximately equal. Insert the 8 stripped fibers through an SMA strain relief shroud, sleeve and SMA connector, then place tips under stereo microscope. Align the 8 stripped fiber tips against a small tolerance flat machined metal part (such as a Thor Labs mount base BA1) so the 8 fibers are aligned to form a plane. Apply just enough optical adhesive to coat and adjoin the fibers. Cure with UV light and eye protection.
6. Next step is to form the line of fibers that matches the slit shaped aperture on the spectrometer. Place an SMA connector in a clamp under stereoscope and attach the fiber bundle such that its position can be adjusted using a micrometer driven translation stage (such as Thor Labs PT1). Adjust the fiber bundle using the translation stage until the line of fibers aligns with the center line of the SMA connector tip (the fiber *bundle* should align with the central longitudinal axis of the SMA connector, and the center point of each fiber's circular cross-section should form a line that aligns with the SMA connector's circular aperture centerline). Fix the fiber bundle in the SMA connector using optical adhesive and UV cure. Polish the tip using D50-SMA and polishing films (LF30D-LFCF) with distilled water as in previous polishing steps above. This 8-fiber bundle can be labelled as "detection side."
7. Insert the two cannulae ends into a female 3D printed part against the flat surface of the depth setting jig for Thor Labs polishing set D50-F. Use stereoscope to verify the polished fiber tip faces are flush with the 3D printed part's flat mating surface. Insert magnets similarly, flush with the flat mating surface. Secure the fiber and magnets to opposite side of 3D printed part using epoxy.

**Measurement Instructions: Software Operation**

To use the code with a Windows 10 OS, run App_Installer.exe from the folder SuppMats, which is zipped as a supplementary material to this paper. If no spectrometer is connected, the code can still be run in demonstration mode to fit the data such as in the manuscript Figure 1D.

On the second screen, you must choose a folder with write permission as the destination for installation. The default installation program is in C:\Program Files, but if you leave this default, saving data will fail.

You will see:
Choose installation folder:
C:\Program Files\The_Rockefeller_Univeristy\Cerebral_Spectroscopy
And you need to manually change this text to:

C:\Users**USER_NAME**\Desktop\The_Rockefeller_Univeristy\Cerebral_Spectroscopy

where **USER_NAME** is the user name with which you are logged in. An easy way to determine the path string is to right-click on any file located on your desktop and select properties then in the "Location:" field, you can copy the path to the desktop for installation.

Follow installation instructions and assure the necessary permissions. This will download from the internet and install the necessary Matlab runtime environment, which will be <1GB in size. Once installation is complete, the program "Cerebral_Spectrometer" will be available in the Start Menu and you can, additionally, chose to create a desktop shortcut during installation for convenience.

Upon successful installation, the icon to run the app will be located in:
C:\Users**USER_NAME** \Desktop\The_Rockefeller_Univeristy\Cerebral_Spectroscopy\application

All experimental data will be saved in the following folder:
C:\Users**USER_NAME**\Desktop\The_Rockefeller_Univeristy\Cerebral_Spectroscopy\application\My_Data
where “My_Data” is the folder automatically created to save data in and named according to the string you can edit in the GUI_App, which has the default string My_Data.

Once running, the first step is to calibrate the probe by aiming the fiber at a white reflector such as a Spectralon 99% diffuse reflectance standard and pressing the calibrate button. This will store the calibration spectrum. Then, with magnetic coupling to a brain-fixed probe tip, the second button will acquire and process spectra for the number of seconds that the user has input and save to a folder that will be automatically created with the name specified in the user text field to store the data. If no spectrometer is connected, the code can still be run in demonstration mode to fit the data such as in the manuscript Figure 1D.

Open-source Matlab editing: Using Matlab to modify this program is possible through the App Designer by running the App Designer command on the file included “GUI_App.mlapp”

- Set the current directory in the Matlab environment to the folder of the unzipped supplementary material folder.
- Type in the command window: “appdesigner GUI_App” to edit the open source code and/or modify the graphic user interface layout.

Depreciated GUIDE Matlab command usage: Using Matlab to modify this program is possible through the GUIDE command to modify the user interface and the correlating code GUI.m Execute GUI.m in the MATLAB computing environment.

The spectrometer must be controlled using the OmniDriver java classes directly by adding the OmniDriver path to the text file titled “Driver_Directory_Name.txt” The default value for this path is:
C:\Program Files\Ocean Optics\OmniDriver\OOI_HOME\OmniDriver.jar
This default value may not need adjustment, provided that installation of OmniDriver was in the path specified by the text file.
